# *Aedes aegypti* eggs use rewired polyamine and lipid metabolism to survive extreme desiccation

**DOI:** 10.1101/2022.12.30.522323

**Authors:** Anjana Prasad, Sreesa Sreedharan, Baskar Bakthavachalu, Sunil Laxman

## Abstract

Upon extreme water loss, some organisms pause their life cycles and escape death, in a process called anhydrobiosis. While widespread in microbes, this is uncommon in animals. Mosquitoes of the *Aedes* genus are vectors for several viral diseases in humans. These mosquitoes lay eggs that survive extreme desiccation and this property greatly enhances geographical expansion of these insects. The molecular principles of egg survival and hatching post-desiccation in these insects remain obscure. In this report, we find that eggs of *Aedes aegypti*, in contrast to those of *Anopheles stephensi*, are true anhydrobiotes. *Aedes* embryos acquire desiccation tolerance at a late developmental stage. We uncover unique proteome-level changes in *Aedes* embryos during desiccation. These changes reflect a metabolic state with reduced central carbon metabolism, and precise rewiring towards polyamine production, altered lipid levels and enhanced lipid utilization for energy. Using inhibitor-based approaches targeting these processes in blood-fed mosquitoes that lay eggs, we infer a two-step process of anhydrobiosis in *Aedes* eggs, where polyamine accumulation as well as lipid breakdown confer desiccation tolerance, and rapid lipid breakdown fuels energetic requirements enabling the revival of mosquito larvae post rehydration.

## Introduction

Life as we know has evolved with water. The fundamental unit of life, cells, are made up of water, inorganic ions, and organic compounds. Of these, water makes up a bulk of the cell volume and mass (Cooper & Hausman, 2007; Milo & Phillips, 2015). Water is an active constituent of cells both structurally and as a polar, amphoteric reagent (Ball, 2008, 2017). The versatility and adaptability of water enables important chemical reactions within a cell, maintaining their structure and function (Nelson & Cox, 2004). This makes water vital for life and given this, it is remarkable that some organisms, called anhydrobiotes, can survive extreme water loss (Alpert, 2005; Crowe, 2014; Leprince & Buitink, 2015). The phenomenon of survival after desiccation is common in unicellular microbes, is selectively observed in some plants, rotifers, nematodes, larvae of certain insects, and is seldom found in other organisms (Alpert, 2006). The loss of water, and the associated volume reduction leads to the destruction of cell components, severe mechanical stress, DNA and RNA damage, redox imbalances that lead to oxidative stress, protein denaturation, and the formation of toxic aggregates (Leprince & Buitink, 2015). In general, a cell must protect its integrity, preserve protein function, and protect its genome to survive water loss. Our current understanding of cellular and molecular processes that enable desiccation tolerance in some organisms comes from a limited number of model systems. From organisms like yeasts, tardigrades and nematodes, the following mechanisms have emerged, which suggest processes by which cellular integrity and function can be preserved. Some organisms accumulate the disaccharide trehalose, which replaces water and prevents protein denaturation and changes in membrane conformation (Calahan *et al*., 2011; Erkut *et al*., 2011, 2013, 2016; Tapia *et al*., 2015; Tapia & Koshland, 2014). Other organisms induce intrinsically disordered or chaperone proteins that protect other proteins from denaturation, as well as against damage due to oxidative stress (Boothby *et al*., 2017; Hesgrove & Boothby, 2020). Yet other induced processes in anhydrobiosis include polyamine biosynthesis, xenobiotic detoxification and lipid metabolism (Erkut *et al*., 2013). All these suggest processes that help protect a cell during water loss. Understanding the mechanisms through which some cells and organisms escape death due to desiccation is therefore of fundamental importance, with important implications for agriculture, pest control, and regenerative biology.

Only a handful of insects exhibit desiccation tolerance (Thorat & Nath, 2018). Mosquitoes are vectors for numerous diseases, and are quintessential examples of insects that require water to breed. *Aedes* mosquitoes lay eggs in fresh water, which is required for the larvae to hatch. Paradoxically, the *Aedes* genus of mosquitoes, responsible for a host of arboviral diseases including dengue, Zika, yellow fever and Chikungunya, lay eggs that tolerate desiccation and can remain in a dormant state for prolonged periods of time – year or more (Mayilsamy, 2019). This phenomenon is reversible and eggs hatch immediately into larvae upon contact with water (Faull *et al*., 2016; Mayilsamy, 2019). By using such strategies, the *Aedes* mosquitoes have globally expanded beyond their original habitats in sub-tropical north Africa (Diniz *et al*., 2017; Kraemer *et al*., 2015; Powell & Tabachnick, 2013).

Currently, the mechanisms associated with desiccation tolerance or survival after rehydration in *Aedes* eggs are unknown. Here, we address the biochemical changes in eggs of *Ae. aegypti* that enable tolerance to extended periods of desiccation and subsequent hatching of eggs after rehydration. We uncover both general, as well as unique biochemical underpinnings in the *Aedes* mosquito egg for desiccation tolerance. These findings provide a biochemical basis for desiccation tolerance in a major vector of arboviral diseases.

## Results and discussion

### *Aedes* eggs acquire desiccation tolerance during late embryonic development

Field studies with *Ae. aegypti* found that *Aedes* eggs are resistant to desiccation, survive for months in a dormant state, and hatch into first instar larvae when submerged in water (Mayilsamy, 2019). To systematically understand this process, we developed a controlled, quantifiable desiccation and hatching assay for *Aedes* eggs (Figure 1A). Eggs of *Ae. aegypti*, along with eggs from a distinct mosquito species *Anopheles stephensi*, were dried for up to 21 days (see Materials and methods). These eggs were revived by placing them in water, and the number of larvae that hatched were counted (Figure 1A). Eggs from the 0-day batch that were not subjected to drying (fresh eggs) were observed under a stereomicroscope, and these eggs appeared intact and healthy. In contrast, after 21 days of storage and desiccation, desiccated eggs appeared deformed with inward shrinkage (Figure 1B). These eggs had a ∼65% reduction in total mass (Figure 1B). Notably, the *Aedes* eggs showed excellent viability upon rehydration. When 21-days desiccated *Aedes* eggs were rehydrated, ∼85% of the eggs rapidly hatched into viable larvae (Figure 1C). The eggs from all the batches tested hatched within 30 minutes of placing them in water. In contrast, *An. stephensi* eggs did not resist even mild desiccation, and no eggs hatched even after 3 days of desiccation (Figure 1C). This establishes *Ae. aegypti* eggs as true anhydrobiotes.

**Figure 1.**
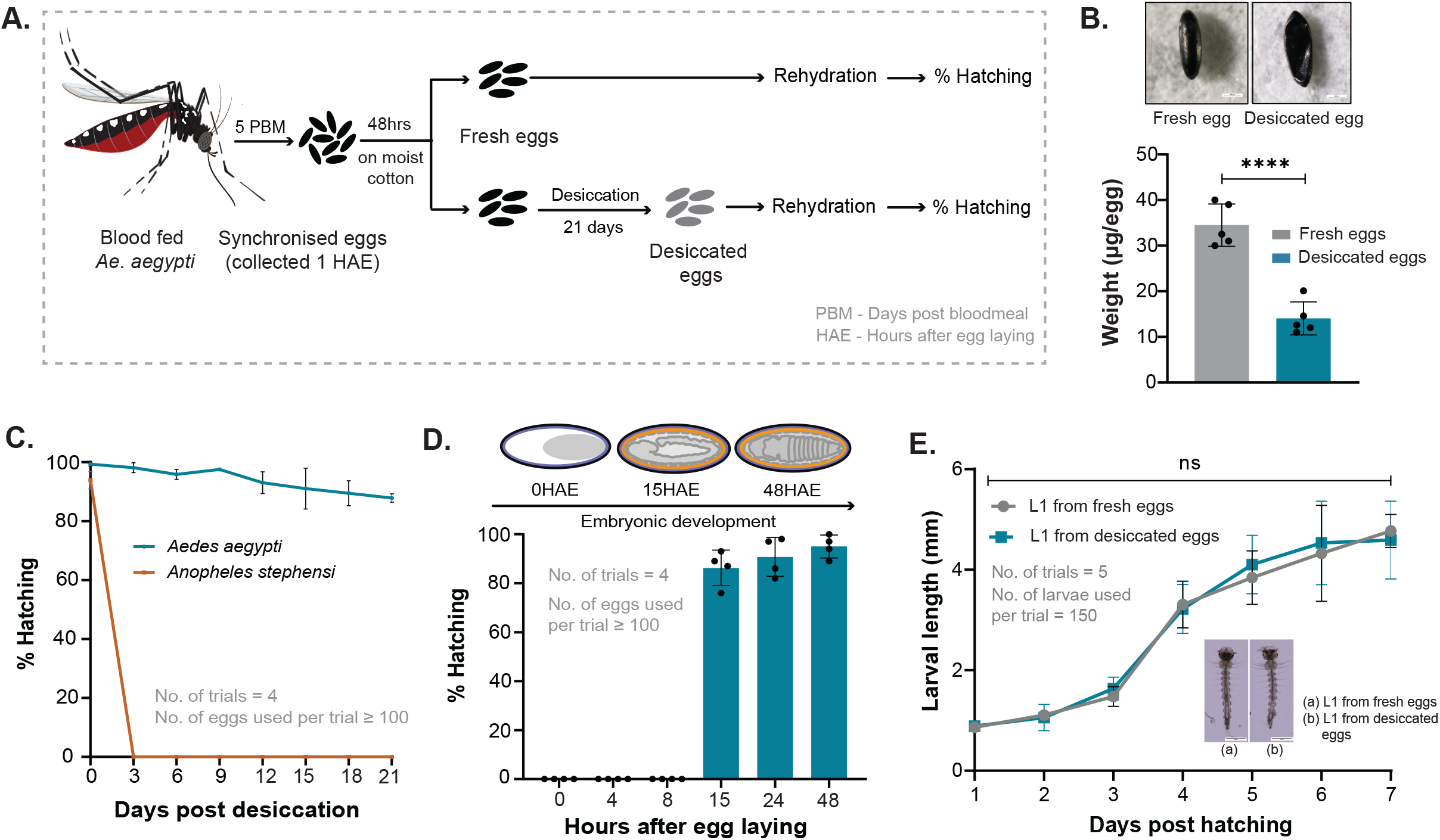
*Aedes* eggs acquire desiccation tolerance during late embryonic development. A) Schematic depicting the desiccation assay. Synchronized eggs were collected 5 days post blood meal. Fresh eggs were hydrated and hatched 48 hours after being laid. Desiccated eggs were dried after 48 hours for 21 days. Fresh or desiccated eggs were rehydrated and 1^st^ instar larvae hatching from these eggs was counted to calculate percentage hatching. B) Morphology and weight changes in desiccated eggs. *Aedes* eggs before (top left) and after desiccation (top right). Desiccated eggs appear deformed and shrunk inwards whereas fresh eggs were oval-shaped and healthy. Scale bar = 1000μm. The number of trials = 5. Data is represented as mean ± SD. C) *Aedes* eggs and desiccation survival. The graph shows percentage of desiccated *Aedes* and *Anopheles* eggs hatching over a period of 21 days. Data is represented as mean ± SD. The number of trials = 4, number of eggs used per trial ≥ 100. D) Embryonic development stages and *Aedes* egg resistance to desiccation (ERD). The schematic (top) shows the structure of a mosquito embryo and the egg shell layers over time during embryogenesis (denoted as hours after egg laying - HAE). Egg shell layers in the schematic are represented as: black - exochorion, blue - endochorion, orange - serosal cuticle, grey dotted lines - serosa, grey - developing embryo. The graph shows percentage of desiccated *Aedes* eggs hatching, when collected and dried at different stages of embryonic development. Data is represented as mean ± SD. The number of trials = 4. The number of eggs used per trial ≥ 100. Also see Figure S1A for further analysis of development stage-based hatching. E) Desiccation and larval development. Growth of 1^st^ instar larvae hatching from fresh or desiccated eggs was measured in terms of larval length. Number of trials = 5. Number of larvae used per trial = 150. Inset: Morphology of *Aedes* 1^st^ instar larvae hatching from fresh (left) and desiccated eggs (right). Scale bar = 1000μm. Also see Figure S1B for details on hatching duration. The statistical significance was calculated using an unpaired-student t-test. *p<0.05, **p<0.01, ***p<0.001, ns - no significant difference.

Next, in order to determine if eggs have to develop to a specific stage in order to develop desiccation tolerance, we systematically subjected *Aedes* eggs to desiccation - 4, 8, 15, 24 and 48 hours post egg laying. Eggs at these different stages of embryonic development were desiccated for a total period of ten days, following which they were rehydrated (see Materials and methods). We observed that eggs subjected to desiccation only after 15 hours post egg laying remained viable (Figure 1D). In contrast, all eggs with less than 15 hours of development before desiccation failed to hatch upon rehydration. We therefore draw two conclusions from these data. First, for desiccation tolerance, embryos develop until the formation of the serosal cuticle along with the two eggshell layers, consistent with earlier studies (Farnesi *et al*., 2015; Rezende *et al*., 2008; Vargas *et al*., 2014). However, this alone is insufficient, since the desiccation intolerant *An. stephensi* eggs also have a serosal cuticle. The resistance to desiccation specifically in *Aedes* eggs must therefore come from other factors within these eggs.

### Desiccation does not affect larval development post hatching

Growth of larvae hatching from fresh or desiccated eggs was determined in terms of length of the larvae. Larval length was measured everyday post hatching. There was a steady increase in the lengths, without any significant difference when the hatchlings from fresh eggs were compared to those from desiccated eggs (Figure 1E). There was also no difference in the percentage of pupation and eclosure between hatched fresh eggs and desiccated eggs (Figure S1A). Further, there was no delay in the development as seen from the duration of pupation and eclosure in the larvae/pupae emerging from fresh or desiccated eggs (Figure S1B). We also qualitatively examined total protein in 1^st^ instar larvae after they emerged from fresh or desiccated eggs, after one hour of hatching. Whole protein extracted from these larvae was resolved and visualized on an SDS PAGE gel. At a purely qualitative level, these samples from larvae hatching from fresh or desiccated eggs did not show reproducibly visible differences in their protein profiles (Figure S1C). Collectively, these data indicate that the rehydrated eggs develop into largely normal 1^st^ instar larvae, and that desiccation does not alter post-hatching larval development.

### Desiccated *Aedes* eggs remodel their proteome towards lipid metabolism and TCA cycle

We next asked if during desiccation, the *Aedes* eggs underwent any proteome remodelling. For this we adopted a differential proteome-based approach to identify selective proteome remodelling in desiccated eggs. Whole protein extracted from fresh eggs v/s desiccated eggs was resolved on gradient SDS PAGE gels and stained with Coomassie to visualize proteome-level changes. The protein gel revealed obvious differences in protein profiles (Figure 2A) between fresh and desiccated eggs. Some proteins visibly increased in the eggs of *Aedes* post-desiccation while some decreased. Protein bands from each lane were excised, proteins extracted and identified using mass spectrometric approaches (see Materials and methods).

**Figure 2.**
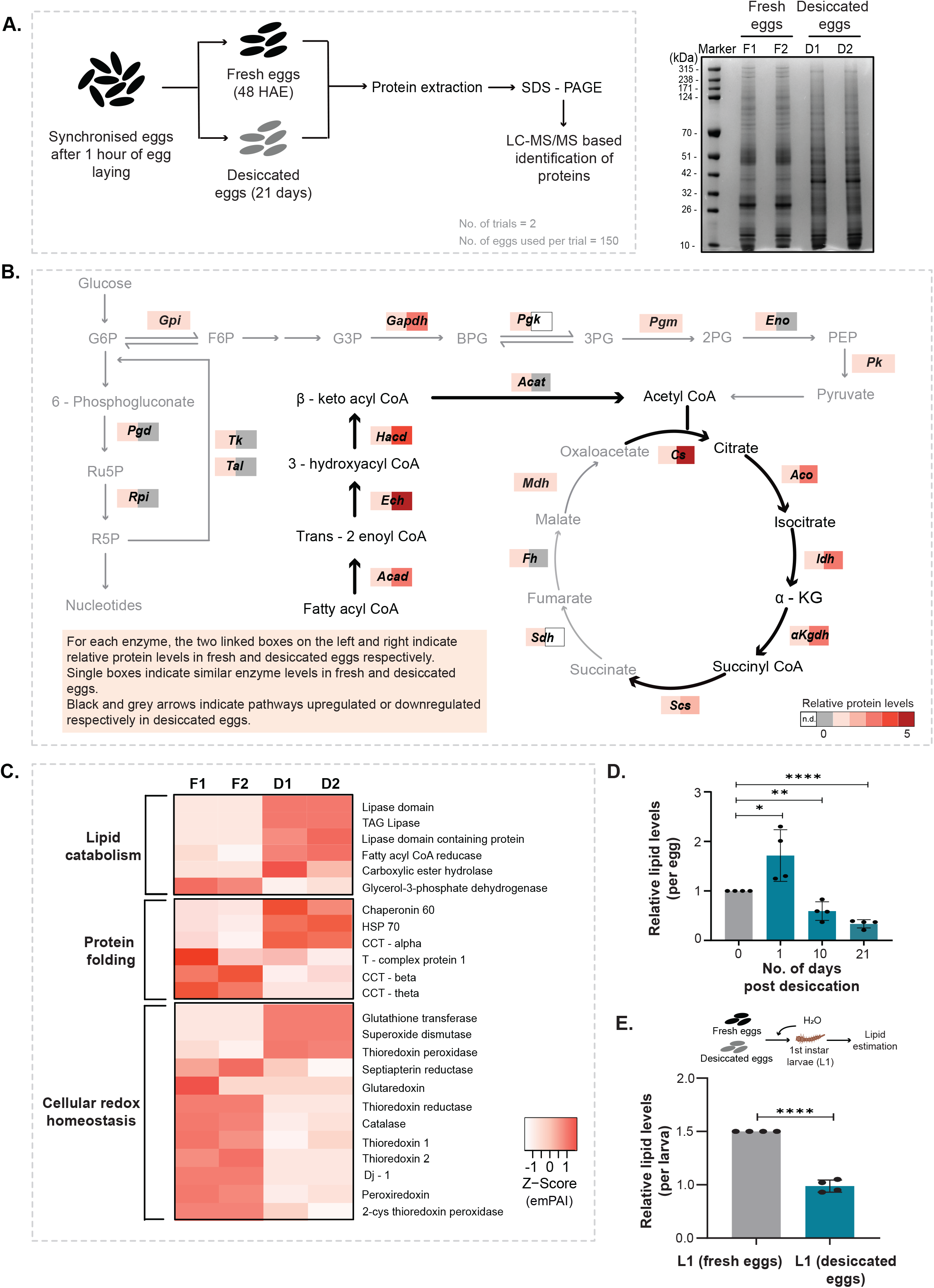
*Aedes* eggs remodel their proteome towards lipid metabolism and TCA cycle. A) Proteome changes in *Aedes* eggs post desiccation. (Left) Schematic showing the experimental setup. (Right) Coomassie stained SDS-PAGE gel of fresh (F1 and F2) and desiccated eggs (D1 and D2), which show a clear difference in band pattern. B) Analysis of proteome changes in fresh and desiccation eggs. The pathway maps of glycolysis, the pentose phosphate pathway (PPP), the TCA cycle, and the β-oxidation of fatty acids are shown, and the colour coded boxes show relative changes in protein levels of enzymes in these pathways based on average emPAI scores (left: fresh eggs, right: desiccated eggs). Note: β-oxidation and the upper arm of the TCA cycle increase in desiccated eggs, while the PPP, glycolysis and the lower arm of the TCA cycle decrease. The number of trials = 2 (F1 and F2 – 2 trials from fresh eggs, D1 and D2 – 2 trials from desiccated eggs). The number of eggs used per trial = 150. Also see Figure S2. C) The heatmap represents differential protein expression in fresh (F1 and F2) v/s desiccated eggs (D1 and D2), for key proteins from three groups: lipid metabolism, protein folding and redox homeostasis. The colour corresponds to the emPAI score converted to a Z-score for that protein. D) *Aedes* eggs and changes in total lipids upon desiccation. The graph represents relative lipid levels in fresh eggs desiccated for 1, 10 and 21 days. Data is represented as mean ± SD. The number of trials = 4. The number of eggs used per trial = 50. E) Total lipid levels in 1st instars hatching from desiccated eggs. Schematic at the top shows the experimental setup. The graph represents relative lipid levels in 1st instar larvae hatching from fresh eggs and those hatching from 21 days desiccated eggs. Data is represented as mean ± SD. The number of trials = 4. The number of larvae used per trial = 50. The statistical significance was calculated using an unpaired-student t-test. *p<0.05, **p<0.01, ***p<0.001, ns - no significant difference.

We further analysed MS data to identify proteins and pathways that were uniquely induced upon desiccation. A total of 2141 and 1837 proteins were identified in fresh *Aedes* eggs (replicate 1 and 2 respectively), and 1802 and 1777 proteins were identified in desiccated eggs (replicate 1 and 2 respectively). Out of these, only those proteins with a significant peptide match of more than 2, and with significant emPAI (Ishihama *et al*., 2005) scores, were considered for further analysis. Based on the emPAI score, we grouped the data into increased or decreased proteins. Post desiccation, 45 proteins increased and 125 proteins decreased in amounts. The abundance of 30 proteins did not change during desiccation. The unique protein IDs obtained were used to map functional domains using the *Aedes* genome (Matthews *et al*., 2018), and relevant biological processes these proteins are involved in were assigned (see Table S1 for a complete list of proteins increasing or decreasing post-desiccation).

This analysis revealed that the *Aedes* eggs in the desiccated, stress resistant state differed strikingly from fresh eggs at the protein level, with a clear rewiring apparent at the proteome level, as summarized below. Gene ontology (GO) analysis of differentially-expressed proteins revealed significant enrichment of TCA cycle enzymes as partly increased and partly decreased in desiccated eggs (Figure 2B, Figure S2A). There was also an increased abundance of lipid catabolism enzymes which included lipases and fatty acid oxidation enzymes as shown in Figure 2B and Figure 2C. Furthermore, the desiccated *Aedes* eggs also had increased superoxide dismutase, glutathione transferase and thioredoxin peroxidase which affects redox balance (Figure 2C). Studies in other organisms suggest that desiccation results in the production of reactive oxygen species (Cruz de Carvalho, 2008; Contreras-Porcia *et al*., 2011; Erkut *et al*., 2013). Consistent with these observations, *Aedes* eggs also have increased dismutase, catalase or peroxidase levels. This might allow desiccated eggs to manage oxidative stress that might occur during desiccation. Additionally, proteins denature (and therefore aggregate or precipitate) due to water loss, and countering this requires the activity of protective molecular chaperones (Erkut *et al*., 2013). We also observed an increased production of a few protein chaperones that assist in protein folding (Figure 2C). These components of proteome rewiring related to oxidative stress and protein chaperones in *Aedes* eggs appear to be entirely consistent with observations made in other anhydrobiotes (Cruz de Carvalho, 2008; Erkut *et al*., 2013), and appear to be universal strategies to combat desiccation stress.

Notably, a unique proteome-level rewiring of metabolism could be constructed, which could be organized into an underlying, putative hierarchy. The pathway map (Figure 2B) shows relative changes in central-metabolism enzymes from glycolysis, the pentose phosphate pathway, TCA cycle and β-oxidation of fatty acids. The enzymes of glycolysis and pentose phosphate decreased (Figure 2B). In contrast, most proteins for fatty acid oxidation and only the upper arm of the TCA cycle increased. This suggested a precise metabolic rewiring apparent at the proteome level, where ‘growth and anabolism’ related processes of glycolysis and the pentose phosphate pathway (Rashida & Laxman, 2021), as well as the ATP - and NADH - producing arms of the TCA cycle (the post α-ketoglutarate part of the cycle) decreased. In contrast, lipid breakdown, and first three steps leading to the alpha-ketoglutarate arm of the TCA cycle are high. We therefore hypothesized that flux through this top part of the TCA cycle might be high. To address this at the level of the metabolic state of eggs, we first measured lipid levels in fresh and desiccated *Aedes* eggs (Figure 2D). The desiccated eggs had significantly lower amounts of lipids (Figure 2D). To further resolve this observation, we estimated lipids over time, after eggs were subjected to desiccation. We observed increased lipids in eggs subjected to desiccation stress for 1 day, followed by a subsequent decrease (Figure 2D). This suggested that lipids were synthesised by the eggs immediately after sensing desiccation, and were subsequently broken down via fatty acid β-oxidation. We therefore next estimated the levels of lipids in 1^st^ instars hatching from fresh eggs v/s those hatching from 21 days old eggs. Consistently, we observed reduced levels of lipids in larvae hatching from desiccated eggs (Figure 2E).

The overall proteome rewiring in mosquito eggs therefore suggests a primarily metabolic rewiring. In two other anhydrobiotes, the dauer stage of the nematode *C. elegans*, and in tardigrades, desiccation increases the amounts of intrinsically disordered proteins or IDPs (Boothby *et al*., 2017; Erkut *et al*., 2013), and these proteins are thought to provide protection analogous to protein chaperones. In contrast, in desiccated *Aedes* eggs, there is no induction of proteins that might fall into this category of IDPs, nor are there any orthologs of the tardigrade or nematode IDP proteins present in the *Aedes* genome. We therefore minimize the possibility of induced IDPs being a mechanism of desiccation tolerance in mosquito eggs, and hypothesize a primarily metabolic rewiring-based acquisition of desiccation tolerance in these eggs.

### Desiccated eggs acquire a hypometabolic state with increased polyamine accumulation

Based on the distinct proteome remodelling after desiccation we constructed a putative, hypothetical metabolic program in the *Aedes* egg correlated with desiccation tolerance. We further asked if this remodelled metabolic state in desiccated *Aedes* eggs retained features of other anhydrobiotes. A feature of anhydrobiotes is to alter carbon metabolism to what has been described as an ‘ametabolic state’ (Erkut *et al*., 2016; Ryabova *et al*., 2020), but this should be more correctly described as a rewired metabolic state with reduced ATP-generating glycolysis and TCA cycle output. Desiccation induces oxidative stress and increased generation of reactive oxygen species (ROS) which leads to reduced cell viability (Contreras-Porcia *et al*., 2011; Rapoport *et al*., 2019). Studies in multiple organisms find a downregulation of the TCA cycle (Diniz *et al*., 2017; Erkut *et al*., 2016; Hell *et al*., 2019; Poelchau *et al*., 2013; Ryabova *et al*., 2020; Zhang *et al*., 2019). This reduces ATP production, but will concurrently also reduce the production of NADH and ROS from the electron transport chain (ETC). Our proteomic data suggested a substantial decrease in glycolysis as well as the lower arm of the TCA cycle. To directly assess this, we first estimated steady-state levels of key glycolytic intermediates and pentose phosphate pathway intermediates. The amounts of these intermediates were either reduced or remained constant in desiccated eggs (Figure 3A, Figure S3A). This is consistent with the proteomic data, and indicates that these eggs have reduced glycolysis. An added outcome of reduced glycolytic metabolism can in some cases be an increase in trehalose synthesis, due to ‘overflow’ metabolism (Erkut *et al*., 2011, 2016; Gupta *et al*., 2019). Trehalose is a versatile desiccant, and in nematodes and yeasts, carbon flux towards trehalose synthesis increases during desiccation (Calahan *et al*., 2011; Erkut *et al*., 2016; Tapia & Koshland, 2014). To assess if this occurred in *Aedes* eggs, we estimated trehalose amounts in fresh and desiccated eggs (Figure S3B). Notably, in *Aedes* eggs, the amounts of trehalose were very low, and did not increase post desiccation (Figure S3D). Note: Controls that included trehalose estimates from comparable biomass of budding yeast grown in glucose in log-phase, which at this stage are themselves not desiccation resistant (Erkut *et al*., 2016), had an order of magnitude greater amounts of trehalose than the *Aedes* eggs (Figure S3D). These data diminish the possibility of protective roles of trehalose in *Aedes* eggs during desiccation.

**Figure 3.**
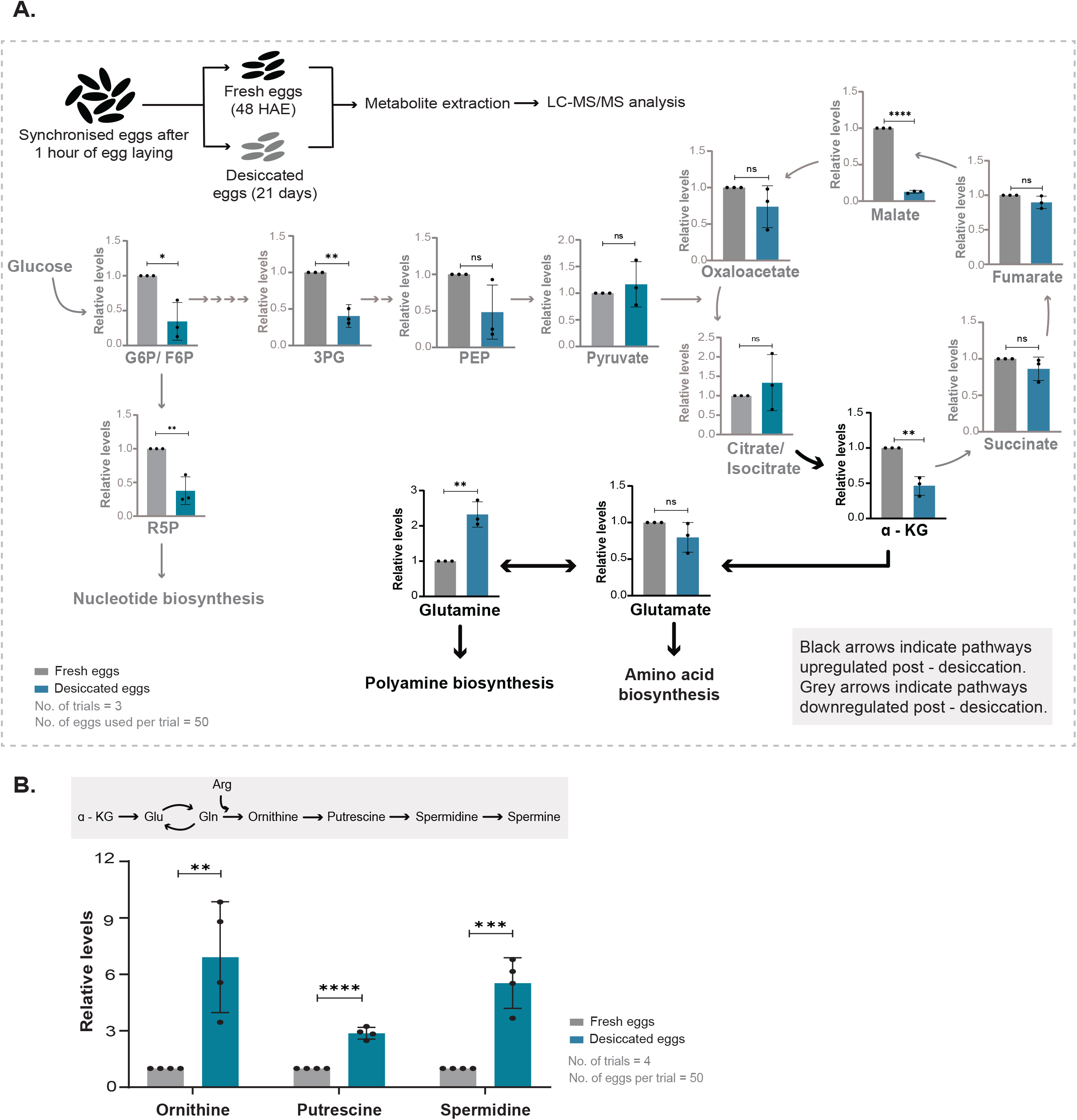
Desiccated *Aedes* eggs acquire a hypometabolic state with increased polyamines accumulation. A) Top: Schematic showing the experimental set up for metabolite extraction and estimation of steady state metabolites in respective pathways. Bottom: The graph represents relative steady state levels of specific amino acids, glycolysis and TCA cycle metabolites. G6P – glucose-6-phosphate, F6P – fructose-6-phosphate, 3PG – 3-phosphoglycerate, PEP – phosphoenolpyruvate, R5P – ribose-5-phosphate, α-KG – alpha ketoglutarate. Data is represented as mean ± SD. The number trials = 3. The number of eggs used per trial = 50. Also see Figure S3A, S3B and S3C for other measured sugar phosphates, trehalose amounts and amino acids. B) Accumulation of polyamines in desiccated eggs. (Top) Pathway of polyamine synthesis, derived from the upper arm of the TCA cycle and arginine metabolism. (Bottom) The graph represents relative amounts of ornithine, putrescine and spermidine. Data is represented as mean ± SD. The number trials = 3. The number of eggs used per trial = 50. Also see Figure S3D for data from *An. stephensi*. The statistical significance was calculated using an unpaired-student t-test. *p<0.05, **p<0.01, ***p<0.001, ns – no significant difference.

We next assessed amounts of the TCA cycle and related metabolites (Figure 3A). Here, we observed a small increase in citrate/isocitrate (in the early part of the TCA cycle), but lower TCA metabolites from the later part of the cycle. This would be consistent with a collectively reduced TCA cycle that is coupled with energy production. However, an interesting trend could be observed in that the upper arm of the TCA cycle remained high, also consistent with the proteomics data. This could suggest a more nuanced metabolic rewiring. Notably, the step which leads to the production of α-ketoglutarate is the first step towards glutamate, glutamine, arginine and proline synthesis. Furthermore, arginine is the precursor for polyamine synthesis. We therefore hypothesized that this increased arm of the TCA cycle (to α-ketoglutarate) resulted in a diversion towards these other molecules, while reducing the complete TCA cycle. We therefore first measured amounts of amino acids in fresh and desiccated eggs of *Aedes*. Notably, the levels of glutamine and arginine increased in desiccated eggs (Figure 3A, Figure S3C). We next measured the levels of polyamines in fresh and desiccated eggs. All the polyamines measured – ornithine, putrescine and spermidine increased substantially in desiccated *Aedes* eggs (Figure 3B). As an added comparison, we also estimated polyamine levels in the desiccation incapable *An. stephensi* eggs. Notably, these polyamines decreased in amounts in these eggs (Figure S3D).

Collectively we find that during the process of desiccation there is a shift in metabolism to a hypometabolic state with reduced glycolysis and the TCA cycle, suggesting reduced energy synthesis. Metabolites from the upper arm of the TCA cycle contribute to the production of amino acids, particularly arginine. Subsequently, polyamines, which are derived from arginine metabolism, substantially accumulate during desiccation.

### Polyamines are essential for desiccation tolerance of *Ae. aegypti* eggs

Studies from dauer stage of *C. elegans* larvae found that the polyamine producing-enzymes ornithine decarboxylase and spermidine synthase increased during dauer larvae desiccation (Erkut *et al*., 2013), suggesting that polyamines might enable desiccation tolerance. In general, polyamines have pleiotropic ‘protective’ roles, by binding to nucleic acids, proteins and membrane phospholipids, can also act as ROS scavengers and chemical chaperones, and form liquid crystals, all of which are useful for desiccation tolerance (Miller-Fleming *et al*., 2015; Saminathan *et al*., 2002).

Since the desiccated *Aedes* eggs specifically accumulated polyamines, we hypothesised that polyamines might assist in desiccation tolerance. Since polyamines are essential for life (Miller-Fleming et al., 2015), we had to adopt a titrated, inhibitor-based experimental set-up as best possible, to test their importance in desiccation tolerance (Figure S4A). We used difluoromethylornithine (DFMO) to inhibit ornithine decarboxylase, a rate-limiting enzyme in polyamine biosynthesis. We obtained eggs from mosquitoes that were blood-fed with and without DFMO, and subjected these eggs to desiccation, followed by rehydration and assessed viability (Figure S4A). We observed significantly reduced hatching in desiccated eggs obtained from females treated with the inhibitor, while fresh eggs treated with the inhibitor showed a high percentage of hatching (Figure 4A). Consistently, we measured the levels of polyamines in fresh as well as DFMO-treated and untreated eggs, and observed decreased polyamines in eggs that were treated with DFMO (Figure S4B). Note that higher doses of DFMO resulted in poor egg laying in females, and hence this could not be further optimized. Collectively, these data suggest that the accumulation of polyamines is necessary for desiccation tolerance of *Aedes* eggs, and reducing polyamine biosynthesis renders eggs sensitive to desiccation.

**Figure 4.**
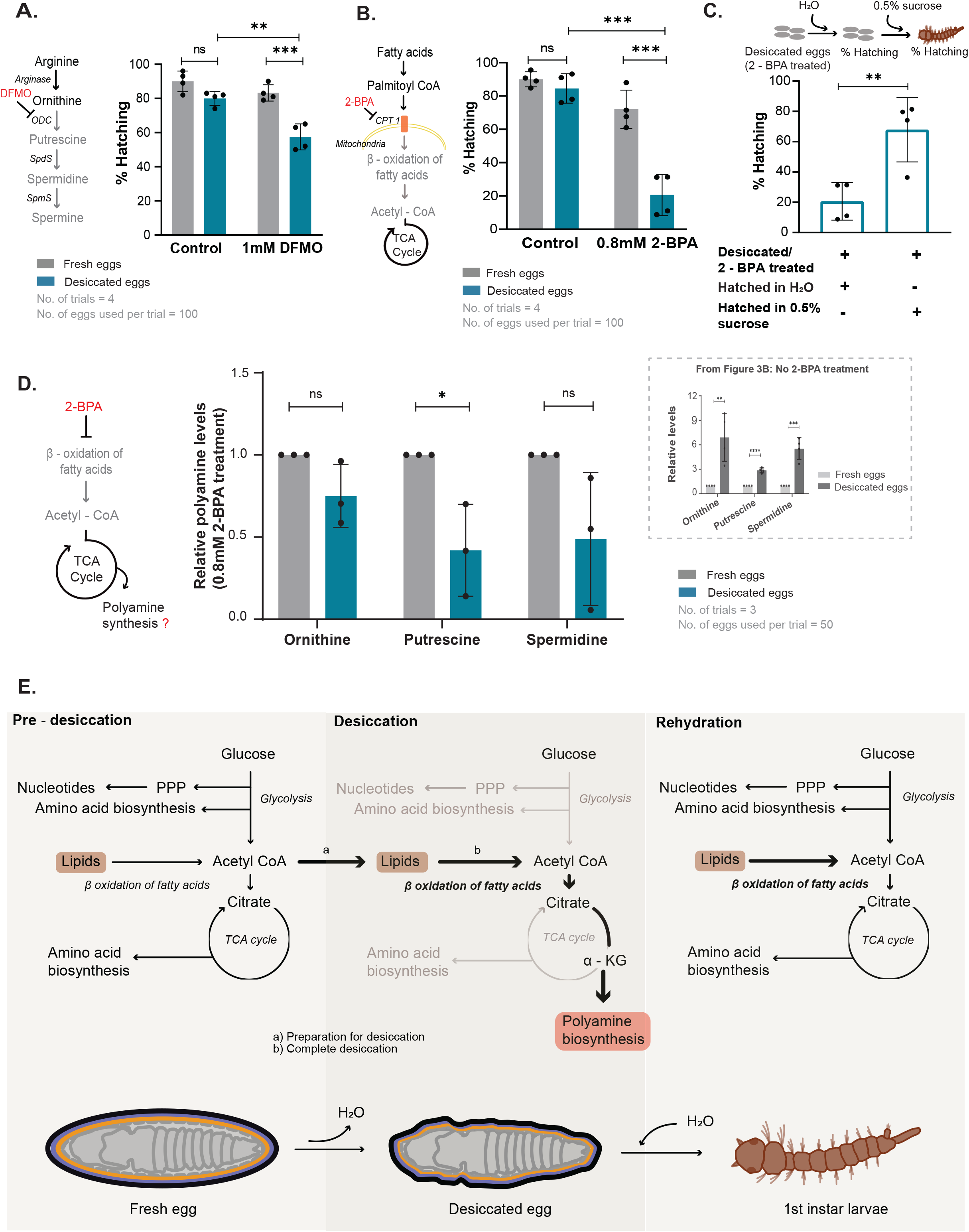
Polyamines and lipid breakdown function synergistically to enable *Aedes* egg desiccation tolerance and larval revival post-rehydration. A) Desiccation tolerance in *Aedes* eggs and dependence on polyamines. Schematic on the left shows inhibition of ornithine decarboxylase (ODC) by difluoromethylornithine (DFMO). Inhibition of the enzyme leads to reduced synthesis of putrescine, spermidine (shown in grey) and accumulation of ornithine (shown in black). The graph shows reduction in the percentage of hatching of eggs post desiccation when treated with DFMO. DFMO - difluoromethylornithine, ODC - ornithine decarboxylase, SpdS - spermidine synthase, SpmS - spermine synthase. Data is represented as mean ± SD. The number trials = 4. The number of eggs used per trial = 100. Also see Figure S4A and S4B for polyamine levels after inhibitor treatment. B) Inhibiting fatty acid oxidation and tolerance to desiccation in *Aedes* eggs. The schematic on the left depicts the mode of action of 2-bromopalmitic acid (2-BPA) which inhibits carnitine palmitoyl transferase 1 (CPT 1), a rate limiting enzyme for β-oxidation of fatty acids in the mitochondria. The graph shows the percentage of fresh and desiccated eggs hatching into 1^st^ instar larvae when females were fed with DMSO (control) and 2-BPA. Data is represented as mean ± SD. The number of trials = 4. The number of eggs used per trial = 100. Also see Figure S4C for lipid levels after inhibitor treatment. C) Requirement for stored lipids in recovery upon rehydration. The top schematic shows the experimental set up, where hatching of desiccated mosquito eggs treated with 2-BPA and hatched in water, was compared to that of desiccated eggs treated with 2-BPA and hatched in 0.5% sucrose. The graph shows the percentage hatching of eggs to 1st instar. Data is represented as mean ± SD. The number of trials = 4. D) Requirement of lipid breakdown for polyamine levels in desiccated eggs. The graph represents steady-state levels of polyamines - ornithine, putrescine and spermidine in fresh and desiccated eggs under 2-BPA treated conditions. Data is represented as mean ± SD. The number trials = 4. The number of eggs used per trial = 50. The inset within the dashed-line box reproduces Figure 3B, for comparison, representing changes in polyamines upon desiccation, without 2-BPA treatment. Also see Figure S4D for a consolidated schematic of effects of different inhibitions on lipid metabolism. E) Model illustrating the metabolic rewiring in response to complete desiccation in *Aedes* embryos. During desiccation, proteins and metabolites of the main energy producing pathways such as glycolysis and TCA cycle (lower arm) reduce, while there is an increase in polyamine biosynthesis and lipid breakdown. Eggs sense desiccation stress and use lipid metabolism as a strategy to prepare them for the dormant state, including diversion of resources towards polyamine accumulation. These fatty acid reserves are also utilised as an energy source for rapid reactivation of metabolism upon rehydration. The polyamines protect the egg during the dormant state. Thick black arrows indicate pathways upregulated during desiccation and grey arrows indicate the pathways downregulated during desiccation.

### Polyamine synthesis and lipid breakdown function synergistically to enable *Aedes* egg desiccation tolerance and larval hatching upon rehydration

Next, we assessed the importance (for desiccation and revival) of the observed increase in fatty acid breakdown pathways, also correlating with altered lipid reserves in desiccated *Aedes* eggs. The need for fatty acid breakdown for desiccation tolerance was not immediately apparent. We considered two scenarios: one where fatty acid breakdown was critical for desiccation tolerance, and a second, more nuanced possibility where fatty acid breakdown enabled survival post rehydration by fuelling energy metabolism required to survive post rehydration from an ametabolic state. To investigate the role of fatty acid oxidation in desiccation tolerance we inhibited this pathway using a titrated inhibitor based experimental set up as described earlier (Figure S4A) using 2-bromopalmitate (2-BPA), a carnitine acetyltransferase inhibitor (Figure 4B). Inhibiting lipid oxidation by 2-BPA significantly reduces the viability of desiccated eggs, as observed by a sharp decline in hatching (Figure 4B). In contrast, fresh eggs from 2-BPA - treated mosquitoes hatched normally. Since desiccated eggs had lower lipid levels, we further hypothesized that lipid catabolism might serve as a source of energy for rapid reactivation of metabolism upon rehydration. If this were so, when an alternate, excellent energy source is provided to desiccated eggs during rehydration, it should rescue survival. To test this possibility, we used the same experimental system as above, where eggs from mosquitoes treated with 2-BPA were desiccated. However, to these eggs, we provided an alternate, high-energy food source (0.5% sucrose) during rehydration, and then estimated larval survival. When the inhibitor-treated desiccated eggs were rehydrated in the presence of sucrose, we observed near-complete hatching rescue and survival (Figure 4C). These data collectively suggest that a major requirement of increased lipid breakdown in desiccated *Aedes* eggs would be to fuel energy production by providing required precursors for energy metabolism, post rehydration.

Finally, we asked if fatty acid oxidation and lipid catabolism in desiccated eggs was itself coupled to increased polyamine biosynthesis, as part of a comprehensive desiccation-specific metabolic program. The reasoning for this is that polyamine biosynthesis (which comes from α-ketoglutarate and arginine biosynthesis) will require a steady supply of acetyl-CoA. When glycolysis is reduced (as observed in desiccated *Aedes* eggs), lipid breakdown could serve as an alternate source of acetyl-CoA. In such a scenario, inhibiting lipid oxidation should reduce or prevent polyamine accumulation in desiccated eggs. To test this idea, we used 2 - BPA - treated mosquito eggs (in the approach described earlier) and now quantified polyamines in fresh and desiccated *Aedes* eggs. Polyamine amounts were sharply reduced in desiccated eggs treated with 2-BPA (Figure 4D). These data suggested that fatty acid oxidation and polyamine synthesis pathways are coupled in this desiccation program, with fatty acid oxidation also required to maintain the increase of polyamines.

In summary, we uncover a unique, protective metabolic program in *Ae. aegypti* eggs in response to complete desiccation (Figure 4E). In order to survive desiccation, the embryos must first reach an advanced developmental stage only after which their development is arrested as pharate larvae inside the egg, where a sudden drop in humidity is sensed by the embryo to prevent larval hatching. While mosquito eggs have an obvious cuticle that confers some protection to the developing larva, profound metabolic changes within the egg enable resistance to desiccation and survival post rehydration. Upon exposure to drying conditions, the developing embryos alter their proteomes towards a unique metabolic state where ‘energy and growth’ metabolism, along with associated oxidative steps (for example with the later part of the TCA cycle) are reduced. The existing carbon and nitrogen can be channelled towards the production of polyamines, which provides protective functions. Concurrently, the metabolic program shifts towards lipid breakdown which serves two functions. First, lipid breakdown is required to maintain the increased polyamines. Second, lipid breakdown enables embryos to restore energy homeostasis, and refuel recovery once the eggs are rehydrated and the larvae complete their hatching enabling reactivation of larval metabolism after exposure to water. Our study broadly illustrates the organisation of specific metabolic rewiring that collectively protects the developing larva from damage due to water loss, as well as enables recovery and intact hatching when water re-enters the eggs. Given the importance of the *Ae. aegypti* as a primary vector for numerous viral diseases (yellow fever, dengue, chikungunya and others) that affect nearly half the world’s population, as well as the rapid geographical expansion of this mosquito vector, we anticipate that this work will foundationally enable orthologous studies to reduce *Aedes* egg survival and global spread. Additionally, some of the inhibitors described here that reduce desiccation resistance in *Ae. aegypti* eggs, as well as new ones affecting other steps in the egg desiccation pathway, may prove useful as vector-control agents.

## Materials and methods

### Mosquito sources and rearing

*Aedes aegypti* and *Anopheles stephensi* were maintained at the insectary at DBT-inStem and the Tata Institute for Genetics and Society. Adults were fed with 0.2% methyl paraben, 8% sucrose, 2% glucose and a multivitamin solution (Polybion SF Complete, Merck Ltd). 10-day old females were blood-fed using the Hemotek membrane feeding system. Larvae were reared in rectangular trays and fed on a liquid diet mixture of dog biscuits and yeast. Appropriate approvals from the institutional biosafety committee of DBT-inStem were obtained.

### Synchronous egg laying

Female mosquitoes lay eggs 3-4 days post a blood meal. 10 gravid females were chosen randomly from cages and transferred to plugged tubes containing moist cotton and lined with moist filter paper. Tubes were placed in a humid, light-protected chamber at 28°C. In all cases oviposition lasted for 1 hour. The eggs were allowed to complete embryonation by placing the filter papers on moist cotton for 48 hours.

### Determination of desiccation survival time for the eggs of *Aedes* and *Anopheles*

The eggs were divided into eight batches. The egg containing filter papers were desiccated for 21 days. Desiccated eggs were revived at predetermined time points (0, 3, 6, 9, 12, 15, 18 and 21 days) by placing them in trays containing water. The eggs hatch into larvae within 1 day and the larvae are counted. The percentage hatching was calculated as follows:

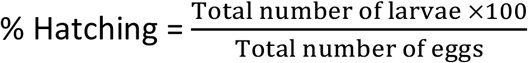

The morphological details of the eggs and the 1st instar larvae were observed under a Nikon SMZ18 stereomicroscope.

### Desiccation assays

Synchronized eggs were collected as described above. Eggs were kept at 27°C on moist cotton until the required age, the onset being considered the end of the 1-hour egg laying period. Embryonic age was assigned as hours after egg laying (HAE). At distinct embryogenesis timepoints (0HAE, 4HAE, 8HAE, 15HAE, 24HAE and 48 HAE) replicates consisting of 100 eggs each, were transferred to a dry Whatman No.1 filter paper and kept under dry conditions for 10 days. Egg viability was assessed by placing each of the filter papers containing eggs in water and counting the 1st instar larvae hatching from it.

### Development of *Ae. aegypti* emerging from desiccated eggs

Larval development was quantified in artificial containers. Synchronous hatching was induced for 60 minutes. 1^st^ instar larvae were placed in trays containing 1.5 liters of RO water. 6 ml of 2% larval food slurry was added for the 1st instars. 2^nd^ instars were fed on 10ml of 2% slurry. For the 3rd and 4th instar 0.5g of larval food was added to 2 liters of RO water. Each tray contained up to 150 larvae. Larval development was quantified in terms of length, as measured daily from the day of hatching till the day of pupation. Performance across conditions was evaluated by measuring the time taken to pupate (days since hatching) and the time to eclose into adults (days since pupation). Percentage hatching, pupation and eclosure were estimated for mosquitoes emerging from fresh or desiccated eggs.

### Proteomics

#### Sample preparation

150 fresh and desiccated eggs of *Aedes aegypt*i were crushed on ice using a pestle in 100μl of 1X phosphate buffered saline (PBS) with 3X protease inhibitor cocktail (Sigma Aldrich, P8340). For protein extraction from 1st instar larvae, synchronous hatching from fresh and desiccated eggs was induced for 1 hour. Larvae hatching were crushed in 1X PBS containing 3X protease inhibitor cocktail (Sigma Aldrich, P8340) supplemented with 10mM PMSF, 10mM iodoacetamide and 4mM EDTA. The samples were sonicated for 5 minutes (5 seconds pulse and 3 seconds rest). The supernatant was collected after centrifuging at 13,000 RPM (10 mins, 4°C) and total protein was estimated by BCA (Pierce BCA Protein Assay Kit, ThermoFisher Scientific, 23225). The supernatant was mixed with 1X Laemmli buffer and boiled at 95°C for 10 minutes. 10ug of each sample was resolved on a 4-12% precast gradient gel (Invitrogen, NP0322BOX), by standard electrophoresis. The gel was stained using Coomassie Brilliant Blue G – 250 (ThermoFisher Scientific, LC6060) for 1 hour, destained, and imaged using the iBright imaging system (ThermoFisher Scientific). Each lane was cut into 4 slices and was separately in gel digested with trypsin, extracted and vacuum dried according to the standard protocol described in (Shevchenko *et al*., 2007).

#### LC-MS/MS

Samples were analysed on Thermo Orbitrap Fusion Tribrid mass spectrometer coupled to a Thermo Nano-flow liquid chromatography system (EASY-nLC 1200 series). Dried digests were reconstituted in 2% acetonitrile/0.1% formic acid and 0.3μg injected onto a LC pre-column (ThermoFisher Scientific Acclaim Pep map 100, 75μm x 2cm, Nanoviper C18, 3μm, 100Å) for separation followed by loading onto the column (ThermoFisher Scientific Easy Spray Pep map, RSLC C18 3μm, 50cm x 75μm, 100Å) at a flow rate of 300nL/min. Solvents used: 0.1% formic acid (Buffer A) and 80% acetonitrile + 0.1% formic acid (Buffer B). Peptides were eluted using a gradient from 10% to 95% of Buffer B for 60 minutes. Full scan MS spectra (from m/z 375 - 1700) were acquired at a resolution of 120000. Precursor ions were sent for subsequent fragmentation by HCD at a collision energy of 32%. MS and MS/MS data were obtained in the orbitrap (Thermo Orbitrap Fusion Tribrid MS, ThermoFisher Scientific).

#### Data analysis

Data analysis was performed using proteome discoverer (version 2.1). The resulting MS/MS data was searched against the *Aedes aegypti* database (Taxonomy id: 7159, 36,032 sequences and 19,407,208 residues). Trypsin was the enzyme used and 2 missed cleavages were allowed. Searches were performed using a peptide mass tolerance of 10 ppm and a product ion tolerance of 0.6 Da resulting in a 1% false discovery rate. A comparison of the identified peptides in desiccated and fresh eggs was done and the data was organised into categories of (i) increased post desiccation (ii) found equally in fresh and desiccated eggs and (iii) decreased post desiccation based on their emPAI scores. Uncharacterised proteins were assessed using PFAM (http://pfam.xfam.org/) for known protein domains and functions were manually assigned. Gene ontology analysis was carried out using VectorBase (https://vectorbase.org/vectorbase/app/). GO terms with corrected p-value <0.05 (Benjamini correction) were considered significantly enriched (see Table S2 for list of all the enriched GO terms).

### Metabolomics

#### Sample preparation

Fifty fresh and desiccated eggs were washed in 80% ethanol (extraction buffer). Eggs were then crushed in 450μl of extraction buffer. Samples were heated for 10 minutes at 85°C and spun at 13,000 rpm for 10 minutes at 4°C. The supernatant was collected into fresh tubes and divided into two parts of 200μl, and one part of 20μl. The parts containing 200μl of the extract were dried using a speed vac.

#### OBHA derivatization

Derivatization in order to detect carboxylic acids was done modifying methods described before (Walvekar *et al*., 2018) using the 20μl part. The derivatized extract was dried using a speed vac.

#### LC-MS/MS and data analysis

Steady state levels of metabolites were analysed using methods described earlier (Walvekar *et al*., 2018). Briefly, extracted metabolites were separated using a Synergi 4-μm Fusion-RP 80 Å (150*4.6 mm, Phenomenex) LC column on Shimadzu Nexera UHPLC system, using 0.1% formic acid in water (Solvent A) and 0.1% formic acid in methanol (Solvent B) for amino acids, nucleotides and TCA metabolites and 5mM ammonium acetate in water (Solvent A) and 100% acetonitrile (Solvent B) for sugar phosphates. The flow parameters were as described (Walvekar *et al*., 2018). Data was acquired using an AB Sciex Qtrap 5500 and analyzed using the Analyst 1.6.2 software (Sciex). Amino acids and TCA intermediates were detected in positive polarity while sugar phosphates were detected in negative polarity mode. The parent and daughter ion *m/z* parameters for metabolites are given in the Table S3. The area under the curve for obtained peaks was obtained using Multi Quant (Version 3.0.1). Analysed data were normalised and plotted (see Table S4 for peak areas of all the detected compounds).

### Trehalose measurements from yeast and mosquito egg samples

10mg of fresh and desiccated *Aedes*’ eggs were transferred to Eppendorf vials. 250μl of 0.25M sodium carbonate was added to all the samples. 10mg of yeast cell pellet (harvested during logarithmic phase of growth) was used for the trehalose assay. 250μl of 0.25M sodium carbonate was added to the cell pellet and all the samples were boiled at 95°C. Enzymatic measurement of trehalose in yeast cells and mosquito eggs was performed according to the protocol described in (Gupta *et al*., 2019; Gupta & Laxman, 2020). Absorbance at 540 nm was determined and compared with the glucose standard to assess the quantity of glucose liberated from trehalose.

### Estimation of total lipids in mosquito eggs and larvae

Total lipids in mosquito eggs and larvae were determined by extraction with 1:1 chloroform methanol followed by a reaction with H_2_SO_4_ and phospho-vanillin reagent as described by (van Handel, 1985). Samples were crushed in 70μl chloroform: methanol (1:1) and the supernatant was used. Lipid standards of 0 - 500μg were prepared such that the final volume was 50μl. The samples were heated at 60°C and the solvents were completely evaporated. 20μl of concentrated H_2_SO_4_ was added to the standards as well as the samples and heated at 100°C for 10 minutes. Samples were brought to room temperature and 480μl of phospho-vanillin reagent was added to the tubes and incubated at 37°C in the dark for 10 minutes for colour development, and absorbance measured at 530 nm.

### Inhibition of β-oxidation of lipids

800μM of 2-bromopalmitic acid (Sigma Aldrich, 238422) was added to fresh blood used for feeding female mosquitoes. Same volume of DMSO was added to blood and fed to another set of females as a control. 5-days post blood meal, females were allowed to lay eggs in ovitraps. Percentage hatching in fresh and desiccated eggs (21-days old) from the control- and inhibitor-treated mosquitoes was calculated. The percentage hatching was further checked 24 hours post revival by replacing water with 0.5% sucrose. Total lipids were estimated in fresh and desiccated eggs obtained from the control and treated mosquitoes using the sulfo-phospho-vanillin method as described above.

### Inhibition of polyamine synthesis in mosquito eggs

1mM of Difluoromethylornithine (Sigma Aldrich, D193) was added to fresh blood and fed to 10-day old females. The same amount of distilled water was added to blood and fed to another set of females as a control. 5 days post blood meal, females were allowed to lay eggs in ovitraps. Percentage hatching of fresh and desiccated eggs (21 days old) from the control- and inhibitor-treated mosquitoes was calculated. Polyamine levels were estimated by mass spectrometry according to the method described above.

### Data visualisation and statistics

All the bar graphs were plotted using GraphPad Prism 8.4.2. Unpaired student’s t-test was used to calculate statistical significance. P-values have been specified in the corresponding figure legends. Heatmap was generated by the software Heatmapper (http://www.heatmapper.ca/). Selected GO terms were visualised in a bubble plot generated using enrichplot package of R and the list of all the enriched GO terms are shown in Table S2.

## Acknowledgements

We thank the NCBS/inStem campus mass spectrometry facility for instrument access and extensive support. We thank Khushboo Agrawal for early help with mosquito rearing and Dr. Sunita Swain (TIGS) for insectary support. We acknowledge extensive support from the insectary at DBT-inStem, and TIGS-CI. BB acknowledges support from Tata Institute for Genetics and Society and SERB (STR/2020/000056). SL acknowledges intramural support from DBT-inStem.

## Figure Legends

**Figure S1: Desiccation and larval development**

A) Desiccation and larval development. The graph shows the percentage of fresh and desiccated eggs hatching into 1^st^ instars, the percentage of larvae developing into pupae and the percentage of pupae developing into adult mosquitoes. Data is represented as mean ± SD. The number of trials = 5. The number of eggs/larvae used per trial = 150.

B) The table shows the duration taken by 1^st^ instar larvae hatching from fresh or desiccated eggs to develop into pupae, and the duration that pupae take to eclose into adults.

C) Whole protein extract from 1^st^ instar larvae hatching from fresh and desiccated eggs analysed on a Coomassie stained SDS PAGE gel. Note: no overt differences in the band pattern in larvae emerging from fresh and desiccated eggs can be observed. The number of trials = 2 (1 and 2 – 2 trials of 1^st^ instar larvae from fresh eggs, 3 and 4 – 2 trials of 1^st^ instar larvae from desiccated eggs). The number of larvae used per trial ∼300.

The statistical significance was calculated using unpaired-student t-test. *p<0.05, **p<0.01, ***p<0.001, ns - no significant difference.

**Figure S2: GO based grouping of proteins that change during desiccation**

A) Gene ontology (GO) based analysis and grouping of proteins into functional categories. The bubble plot shows GO analysis of proteins upregulated in desiccated eggs (black), equally expressed in fresh and desiccated eggs (grey) and proteins downregulated in desiccated eggs (light grey). Rich factor indicated in the y-axis was calculated as a ratio of number of proteins annotated in a particular GO term to total number proteins in that GO term. The colour of each bubble represents the corrected p-values (Benjamini correction) of each term involved in the analysis. The size of each bubble represents the number of proteins identified in this study belonging to the specific GO term.

**Figure S3: Additional metabolite measurements in fresh and desiccated eggs**

A) Steady state levels of other glycolytic in desiccated *Aedes* eggs. The graph represents relative steady state levels of G3P – glyceraldehyde-3-phosphate, F16BP – fructose-1,6 bisphosphate, S7P – sedoheptulose-7-phosphate. Data is represented as mean ± SD. The number trials = 3. The number of eggs used per trial = 50.

B) Trehalose amounts *Aedes* eggs before and after desiccation. The graph represents relative trehalose levels between fresh and desiccated eggs and equal biomass of yeast. The number of trials = 3. Quantity of eggs or yeast used per trial = 10mg.

C) Steady-state levels of all amino acids in fresh and desiccated *Aedes* eggs. The graph represents relative levels of amino acids. Data is represented as mean ± SD. Number trials = 3. Number of eggs used per trial = 50.

D) Polyamine levels in *An. stephensi*, a desiccation sensitive species. The graph represents relative steady state levels of ornithine, putrescine and spermidine. Data is represented as mean ± SD. The number trials = 2. The number of eggs used per trial = 50.

The statistical significance was calculated using an unpaired-student t-test. *p<0.05, **p<0.01, ***p<0.001, ns – no significant difference

**Figure S4: Additional metabolite measurements in inhibitor treated eggs that undergo desiccation**

A) An illustration showing the experimental set up for inhibiting ornithine decarboxylase (ODC) using difluoromethylornithine (DFMO) or fatty acid oxidation using 2-bromopalmitic acid (2-BPA). Mosquitoes were fed with blood containing the inhibitor or the vehicle (H_2_O or DMSO respectively). Desiccation assay as detailed earlier was performed with the fresh and desiccated eggs obtained from the inhibitor fed mosquitoes.

B) Difluoromethylornithine (DFMO) treatment and polyamine levels in desiccated *Aedes* eggs by inhibiting ornithine decarboxylase. The graphs (i-iii) represent steady state levels of polyamines - ornithine, putrescine and spermidine in fresh and desiccated eggs under H_2_O (control) and DFMO treated conditions. Data is represented as mean ± SD. The number trials = 4. The number of eggs used per trial = 50.

C) 2-bromopalmitic acid (2-BPA) treatment for inhibiting beta-oxidation of fatty acids, and lipid levels. The graph represents relative lipid levels in fresh and desiccated eggs under DMSO (control) and 2-BPA-treated conditions. Data is represented as mean ± SD. The number trials = 4. The number of eggs used per trial = 50.

D) Schematic showing the consequences of inhibiting fatty acid oxidation in *Aedes* eggs. Under normal conditions, the stored fats are broken down and feed into the TCA cycle during desiccation and provide energy for them to hatch into 1^st^ instar post-rehydration (a). The percentage of eggs surviving desiccation reduces after fatty acid oxidation inhibition (b). When the eggs are rehydrated in 0.5% sucrose, sucrose serves as an alternate source of energy to sustain the hatching of desiccated eggs (c).

