## Supplementary figures S1 - S4 for "*Aedes aegypti* eggs use rewired polyamine and lipid metabolism to survive extreme desiccation"

### Figure S1: Desiccation and larval development

A.

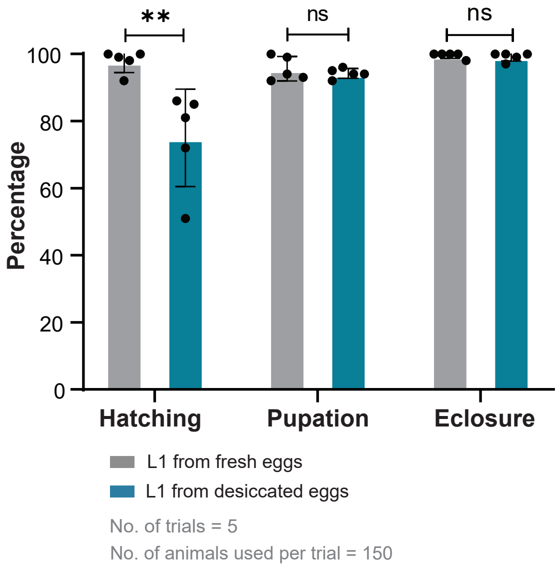


B.

| Duration | L1 from fresh eggs | L1 from desiccated eggs |
|---|---|---|
| L1 to pupa | 7 days | 7 days |
| Pupa to adult | 2 days | 2 days |

C.

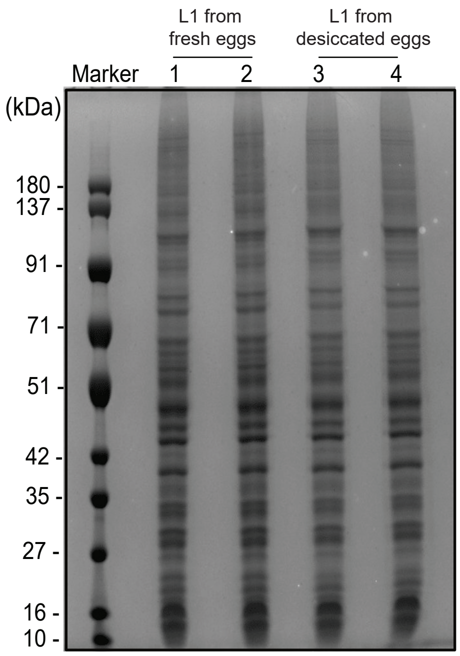

### Figure S2: GO- based grouping of proteins that change during desiccation

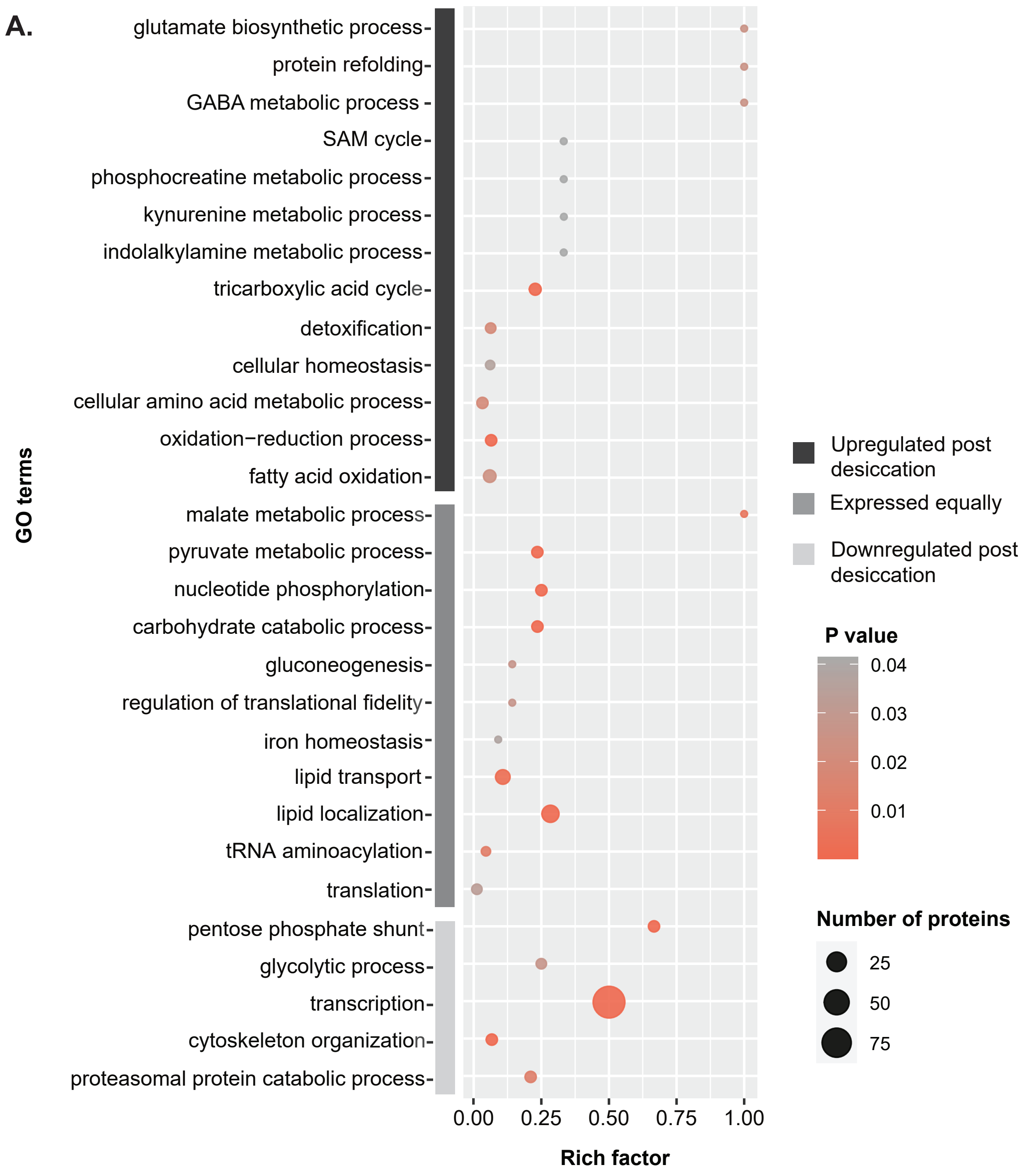

### Figure S3: Additional metabolite measurements in fresh and desiccated eggs

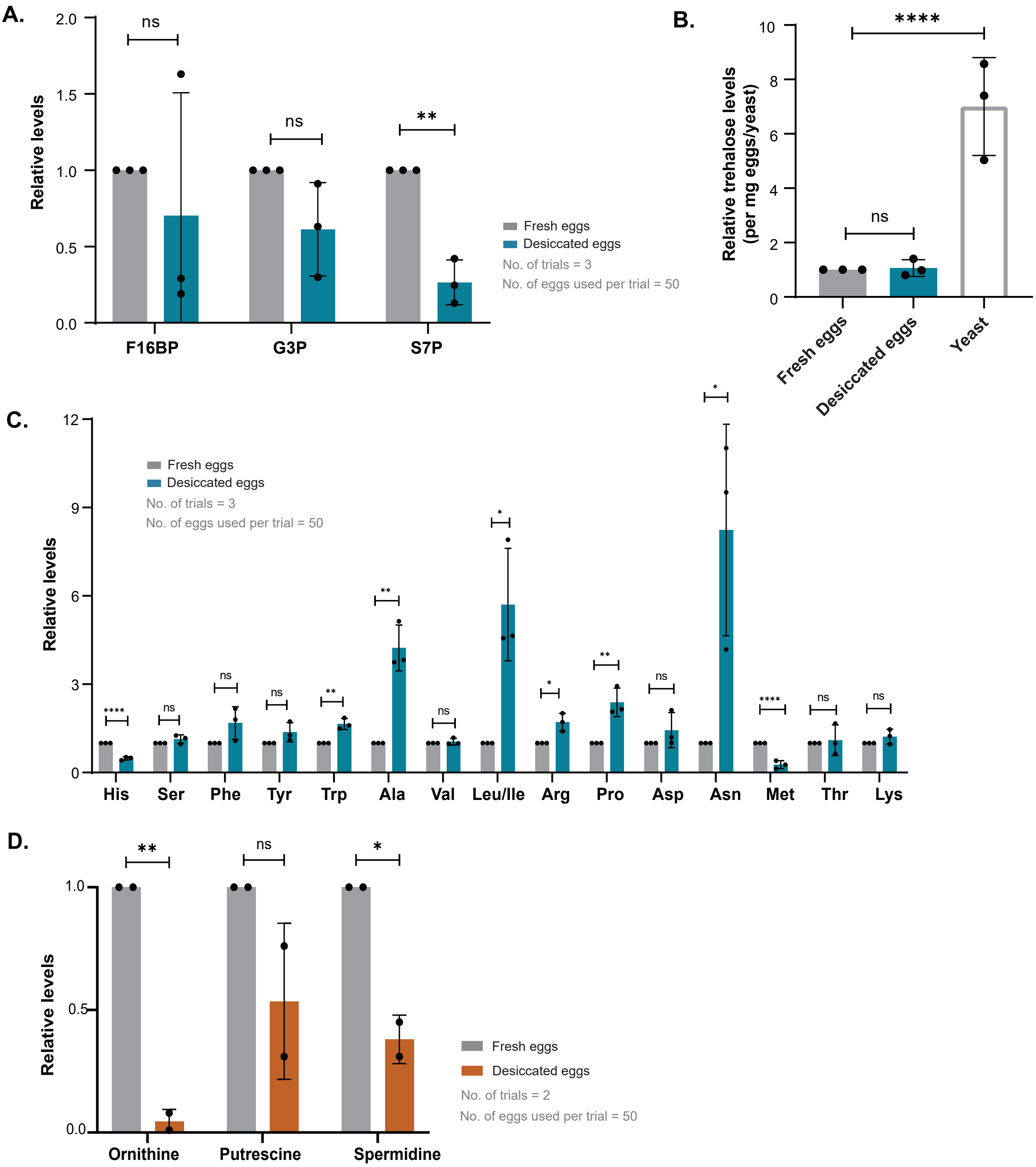

### Figure S4: Additional metabolite measurements in inhibitor-treated eggs that undergo desiccation

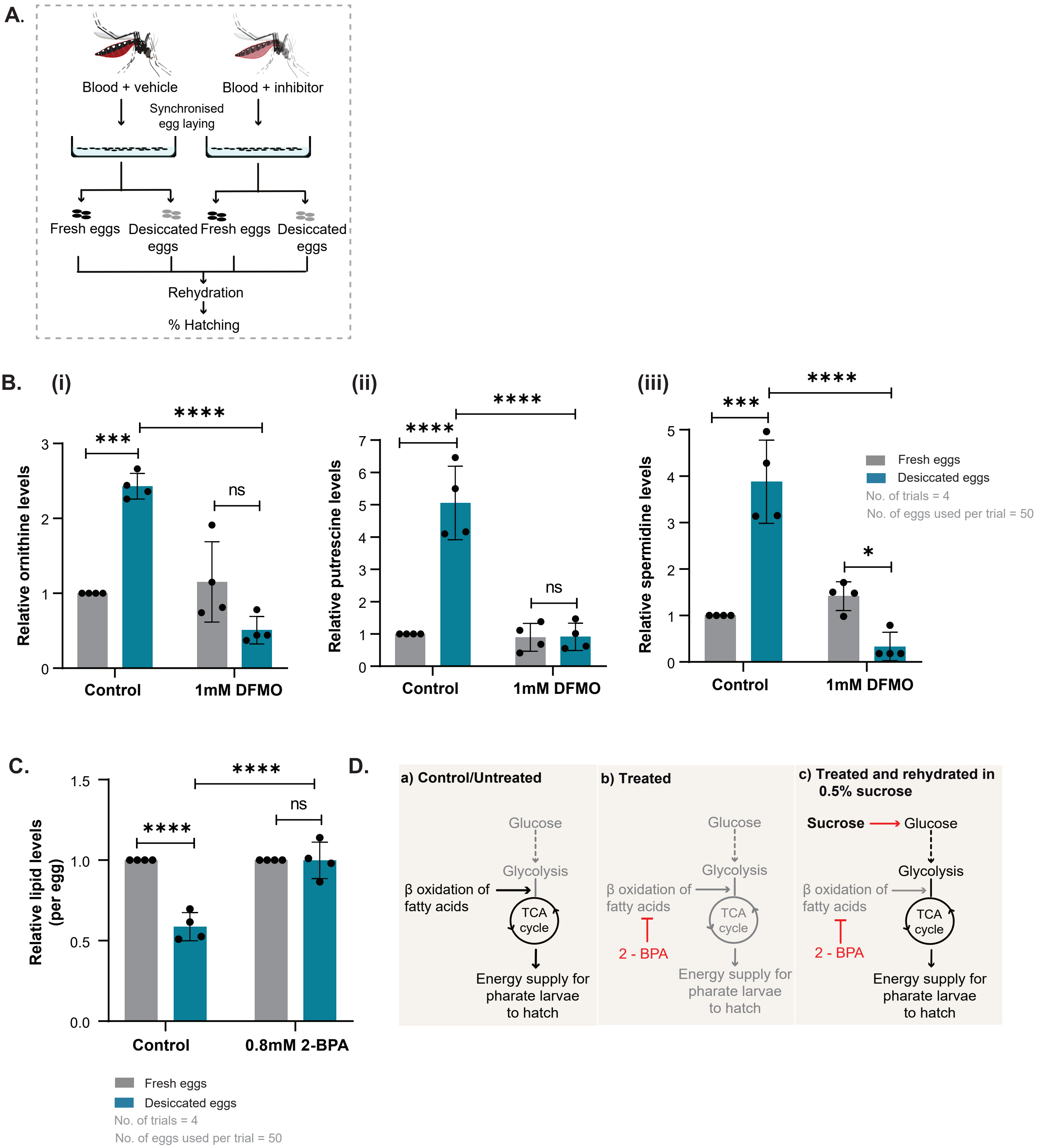
